# Methionine matters: a common mechanism of viral inhibition of host defense identified via AI-assisted molecular dynamics

**DOI:** 10.1101/2025.09.19.677134

**Authors:** Skye J. Bixler, Gregory A. Babbitt, Liv K. Casperson, Maureen C. Ferran

## Abstract

Diverse groups of viruses infecting higher eukaryotes inhibit mRNA export via physical blockage of the Rae1-Nup98 complex within the host cell’s nucleopore. This is thought to most often involve the critical placement of hydrophilic flanked single methionine residues along polypeptide extensions that reach into the nucleopore. However, it is unknown how this presumably conserved mechanism might function across diverse viral taxa. Here we employ a comparative molecular dynamics (MD) approach comparing motions of wild-type and mutant viral proteins in Rae-Nup98 bound vs. unbound states. Our comparisons of MD simulations are enhanced by kernel-based denoising allowing the isolation of non-random functional dynamics from random thermal noise. We demonstrate that despite large structural differences, three evolutionarily distinct viral systems (i.e. VSV M protein, SARS-CoV2 ORF 6, and KSHV ORF 10) share nearly identical single methionine-dependent functional dynamics related to the host cell inhibition of nuclear transport. This finding strongly supports a convergently-evolved common functional mechanism across viruses involving specific structural placement of non-polar residues like methionine and potentially providing a common therapeutic target for broad spectrum anti-viral treatment.

## Introduction

Many viruses employ a common strategy to evade host defenses by inhibiting host mRNA nuclear export. This disruption prevents the expression of key antiviral genes, including interferon-stimulated genes (ISGs), which require export of their transcripts from the nucleus to the cytoplasm for translation (1, 2). Blocking mRNA export not only dampens immune responses early in infection but also redirects cellular translational machinery toward viral protein synthesis, contributing to global host shutoff (3). Additionally, the inhibition of cytokine and chemokine mRNA export delays immune signaling and effector cell recruitment, enhancing viral survival (4).

Despite differences in genome type and replication strategy, Vesicular Stomatitis Virus (VSV), Kaposi’s Sarcoma-Associated Herpesvirus (KSHV), and Severe Acute Respiratory Syndrome Coronavirus (SARS-CoV) target a common host complex—Rae1–Nup98—that is central to the nuclear export factor 1 (NXF1)-mediated mRNA export pathway (2). Rae1 binds single-stranded RNA (ssRNA) through a conserved basic patch and interacts with Nup98 via a GLEBS motif, recruiting NXF1 to facilitate mRNA translocation (5, 6). However, key mechanistic details of Rae1’s function and its viral targeting remain incompletely understood (4, 7, 8).

Each virus expresses a distinct protein to subvert this pathway. Early studies showed VSV M inhibits nuclear export by binding Rae1–Nup98, causing nuclear accumulation of polyadenylated RNA and disrupting nuclear pore components. This protein localizes to the nuclear envelope and shuttles into the nucleus via a nuclear localization signal. Furthermore, a conserved methionine at position 51 is critical as mutation impairs localization and export inhibition (9-12). Structural analyses revealed that the M protein’s N-terminal tail mimics ssRNA, inserting into Rae1’s RNA-binding groove with the conserved methionine anchoring this interaction—mutations disrupt binding and restore export (7, 13).

The ORF6 protein of SARS-CoV and SARS-CoV-2 inhibits mRNA export by binding Rae1–Nup98 through a conserved methionine in their C-terminal tails, blocking antiviral transcript export and interferon signaling. Mutations in this methionine restore export (4, 14, 15). SARS-CoV-2 ORF6 also causes nuclear retention of poly(A)+ RNA, paralleling VSV M’s effect (16). Despite export inhibition, SARS-CoV replication, which is cytoplasmic, remains unaffected (4).

The ORF10 protein of Kaposi’s Sarcoma-Associated Herpesvirus (KSHV), a DNA virus, inhibits host mRNA nuclear export. This viral protein binds directly to the Rae1–Nup98 complex, targeting the conserved RNA-binding groove of Rae1. Unlike RNA viruses that broadly shut down mRNA export, KSHV selectively inhibits export of host transcripts while allowing viral mRNAs to exit the nucleus via the CRM1-dependent pathway. Furthermore, ORF10-mediated inhibition of mRNA nuclear export is impaired by mutations that weaken its interaction with Rae1 (8, 17). This selective inhibition enables KSHV to suppress host antiviral responses without compromising viral gene expression, highlighting a sophisticated viral strategy to balance immune evasion and viral replication (8).

As shown in Supplementary Figure 6, sequence alignments of VSV M, SARS-CoV ORF6, and KSHV ORF10 reveal a conserved methionine residue flanked by acidic amino acids, which enables these viral proteins to mimic host single-stranded RNA and bind Rae1’s positively charged RNA-binding groove (VSV: (9-12), SARS-CoV: (4, 14, 15), KSHV: (8, 17)). However, we note that the large structural diversity in the location of this conserved motif combined with the rapid evolution of viral proteins in general, could also suggest that it arose via convergent evolution. Regardless of its origin, this molecular mimicry competitively inhibits host mRNA export by blocking Rae1’s interaction with cellular transcripts. Functional studies show that substituting this methionine with arginine (M→R) disrupts Rae1 binding and restores normal mRNA export, underscoring the essential role of this residue in viral suppression of host gene expression.

This study uses comparative molecular dynamics simulations to examine how VSV, KSHV, and SARS-CoV viral proteins each interact and bind to Rae1–Nup98 structure. We hypothesize that viral binding induces strong binding interactions that block mRNA export, and that mutating the conserved methionine reverses these binding effects. To test this, we applied the ATOMDANCE software (18), which combines a machine-learning based denoising algorithm with traditional comparative molecular dynamics (MD) simulation to examine subtle functional protein motions at single-site resolution. This denoising can be critical for functional comparisons of molecular dynamics in protein systems where the dynamic motions of important single sites are obscured by large amounts of solvent-induced thermal noise in the majority of the MD simulation. In past studies, we employed very large sampling regimes in order to identify important functional binding sites in MD simulation (19-21). However, the recent addition of kernel-based denoising to our methods allows for identification of single-sites binding effects in as little as 1-10ns of simulation run time (18). In the three viral protein systems, we compare the dynamics of the relevant proteins containing the hydrophilic flanked methionine motif in both the nucleoporin-bound and unbound states. We use an arginine mutation that is experimentally confirmed to prevent Rae1-Nup98 binding in VSV and apply it to each of the critical methionine sites in the three systems and repeat the comparison of nucleoporin bound vs. unbound dynamics in the presence of the destabilizing arginine mutation. We report here on strikingly functional commonalities among structurally diverse viral proteins as well as some important differences in viral binding to Rae1-Nup98 nucleoporin complexes in both wild-type and mutationally destabilized states.

## Methods

### Data Acquisition and Preparation

Crystal structures of the Rae1-Nup98 complex bound to the viral proteins Vesicular Stomatitis Virus (VSV) M protein, Kaposi’s Sarcoma-Associated Herpesvirus (KSHV) ORF6 and SARS CoV-1 & 2 were obtained from the Protein Data Bank (PDB). The viral complexes analyzed included PDB 7BYF, KSHV; PDB 7VPH; PDB 4OWR VSV and 7VPG, SARS CoV-1 & 2 respectively (4, 7, 17). These three viruses were chosen or their proven biological relevance to host mRNA nuclear export inhibition through Rae1 and Nup98, demonstrated in functional assays often involving mutation of hydrophilic/acidic flanked methionine residues in the polypeptide extensions that interact with host Rae1-Nup98 complex in the nucleopore (Figure 1).

**Figure 1.**
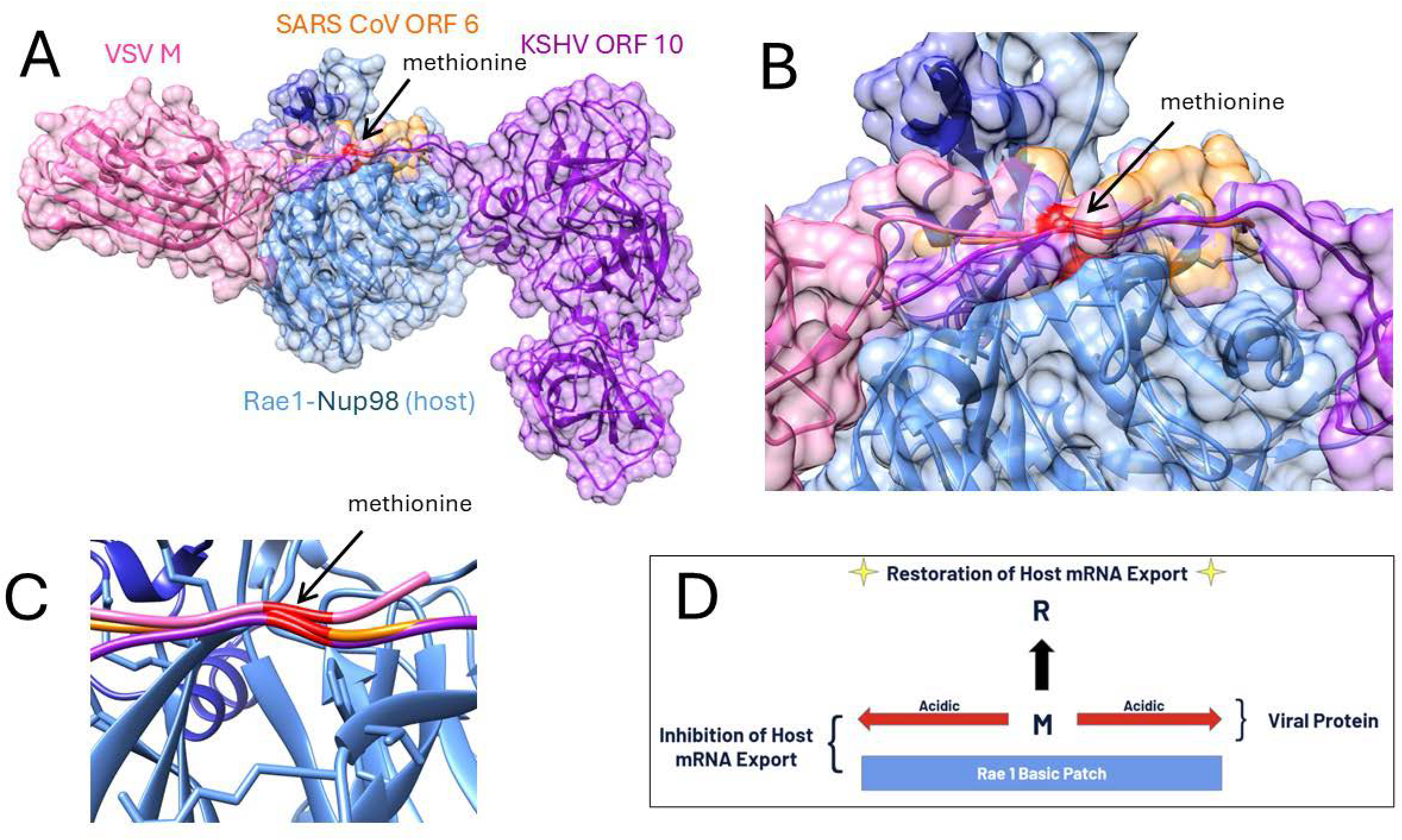
Three viral protein interactions with the Rae1-nup98 complex, (A) a structural overview of the nucleopore penetrating polypeptide extensions of VSV M, KSHV ORF 10, and SARS-CoV ORF 6 in relation to the host target protein complex with (B) close-up view of the functional acidic flanked methionine and (C) a close-up view of the structural overlap of the positioning of the methionine backbone (shown in red). (D) A schematic diagram of the computational experiments performed to release binding and potentially restore host mRNA export function.

Structural preprocessing of the PDB complexes was conducted using UCSF ChimeraX (22). Non-essential molecules within the crystal structures, including water, ions, and ligands, were removed from the six total PDB files.

### Computational Mutagenesis

Mutant versions of each PDB were created using UCSF ChimeraX to introduce a methionine to arginine mutation at position 51 for the VSV M protein, position 58 for the SARS-CoV ORF 6, and position 413 for KSHV ORF10. While on differing proteins and positions, this methionine is shared in its binding grove location within the Rae1-Nup98 complex and is flanked by acidic residues involved in the Nup98-Rae1 binding grove. The following ATOMDANCE workflow was implemented twice on each viral complex to compare the molecular dynamic changes between the viral complex bound and unbound to Rae1-Nup98 as well as to compare wildtype vs a M→R mutant using mutagenesis models created in UCSF ChimeraX.

### Molecular Dynamics Simulations

To prepare the files for molecular simulation, hydrogen atoms were added using the AmberTools package. Energy minimization was performed under the Amber FF14SB force field to ensure structural stability prior to molecular dynamics (MD) simulations (23). All-atom MD simulations were conducted using OpenMM 7.5 (24). Protein systems were solvated in a TIP3P water box with a 10 Å buffer and 0.15 M NaCl added to neutralize the charge. Each system underwent energy minimization followed by NVT and NPT equilibration employing a Langevin thermostat for temperature control and a Berendsen barostat for pressure regulation. Production simulations were run for both one ns and ten ns.

### SITE_WISE Comparison of Dynamic States

The following analytical pipeline within ATOMDANCE software suite (18) was utilized to assess direct site-wise comparisons. DROIDS 5.0 utilized site-wise average differences and Kullback-Leibler (KL) divergences in atom fluctuations. Sites with significantly different dynamics were identified with multiple tests corrected to the sample Komogorov-Smirnov test. Next, a denoised functional comparison was performed with maxDemon 4.0, which utilizes Gaussian kernel process learning and defines the distance between learned features of local atom fluctuation as the maximum mean discrepancy between molecular dynamics of proteins in two functional states.

These two pipelines were utilized for both the wt viral protein bound vs. unbound to the Rae1-Nup98 complex and, secondly, the wild-type vs. M—>R mutant PDBs to quantify the effects of the mutation on molecular dynamics. The comparison between the MD simulations at given sites was reported as maximum mean discrepancy (MMD) in the reproducing kernel Hilbert space (RKHS). Using a bootstrapping approach, hypothesis tests for the significance of functional dynamic changes were also reported via MMD. A key concept here is that because the learning algorithm cannot learn from noise in the atom fluctuations, it can act as a denoising filter and highlight site-wise changes in the non-random or machine-like component of motion. Molecular dynamics simulations are performed to represent the functional end states, such as bound vs. unbound states. The.pdb structure,.prmtop topology, and.nc trajectory files for both states are input into the software, and the trajectories are subsampled using cpptraj as specified by the user. Site-specific local atom fluctuation matrices are generated from the subsampling and used to train a Gaussian process kernel (radial basis function). For each site on the protein, the maximum mean discrepancy (MMD) in reproducing kernel Hilbert space is calculated, quantifying the functional differences in protein dynamics at that site. A positive MMD indicates amplified atom motion, while a negative MMD indicates dampened motion. These MMD values are visualized in plots and can be color-mapped to the.pdb structure in UCSF ChimeraX, with blue indicating regions of reduced motion due to binding interactions and red indicating amplified motion. For more mathematical details of this method please consult our software note in Biophysical Journal (18).

## Results

We observed extensive thermodynamic stabilization, as indicated by large KL divergences in local atom fluctuations, during both 1ns and 10ns simulations of the molecular dynamics of viral protein binding to Rae1-Nup98. We also observed protein-wide destabilization of dynamics due to functional M to R mutations in the polypeptide extensions interacting with target host protein complexes in the nucleopore (Figure 2, Supplemental Figure 1-2).

**Figure 2.**
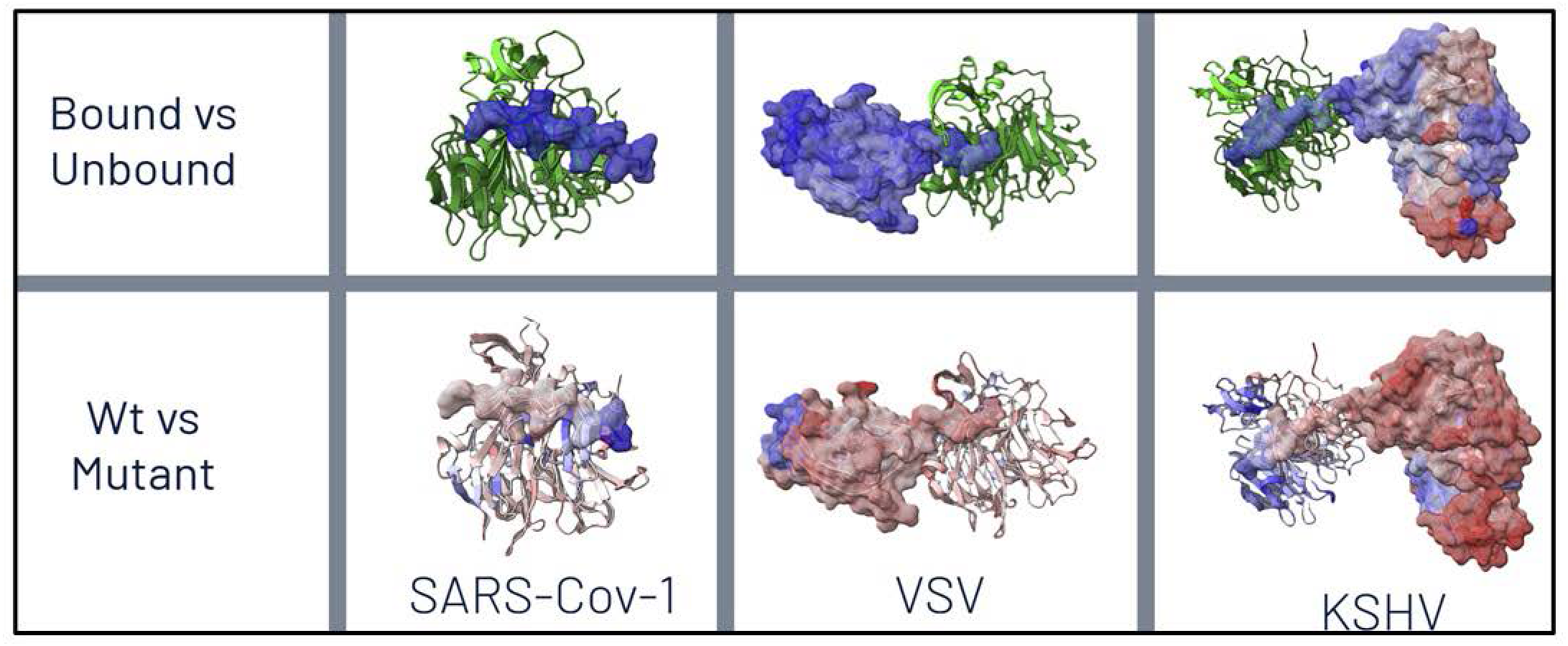
Color-mapping of overall thermodynamic effects of binding (top) and M to R mutation (bottom) on various viral proteins interacting with host Rae1-Nup98 nuceoporin complexes. Here, a signed Kullback-Leibler (KL) divergence in local amino acid backbone root mean square atom fluctuation (rmsf) is shown where the sign imposed upon KL divergence represents dampening of motion or cooling (blue=negative) vs. amplification of motion or heating (red=positive). Here, binding interactions are shown to dampen viral dynamics as expected whereas the M to R mutations in the respective functional sites on the polypeptide extensions are shown to markedly destabilize the binding dynamics of the viral proteins. This effect is very apparent across most of the whole protein in both VSV M and KSHV ORF 10.

In VSV M, these changes were affected a critical single methionine at position 51 as indicated by denoised molecular dynamics comparisons of both binding state and mutational effects when replaced by arginine (Figure 3), a mutation known to restore host mRNA transport in lab strains of VSV (2, 10, 11, 25). A similar mutational effect was shown to be achieved in VSV not only by replacing the methionine at position 51, but also by replacing a key acidic residue in an adjacent site via a D52G mutation (Supplemental Figure 3). While the M51R mutation achieves its destabilizing affects by altering the dynamics of the N-terminal extension of the M protein, the D52G mutation achieved destabilization via release of a loop region in Nup98 (Supplemental Figure 3C).

**Figure 3.**
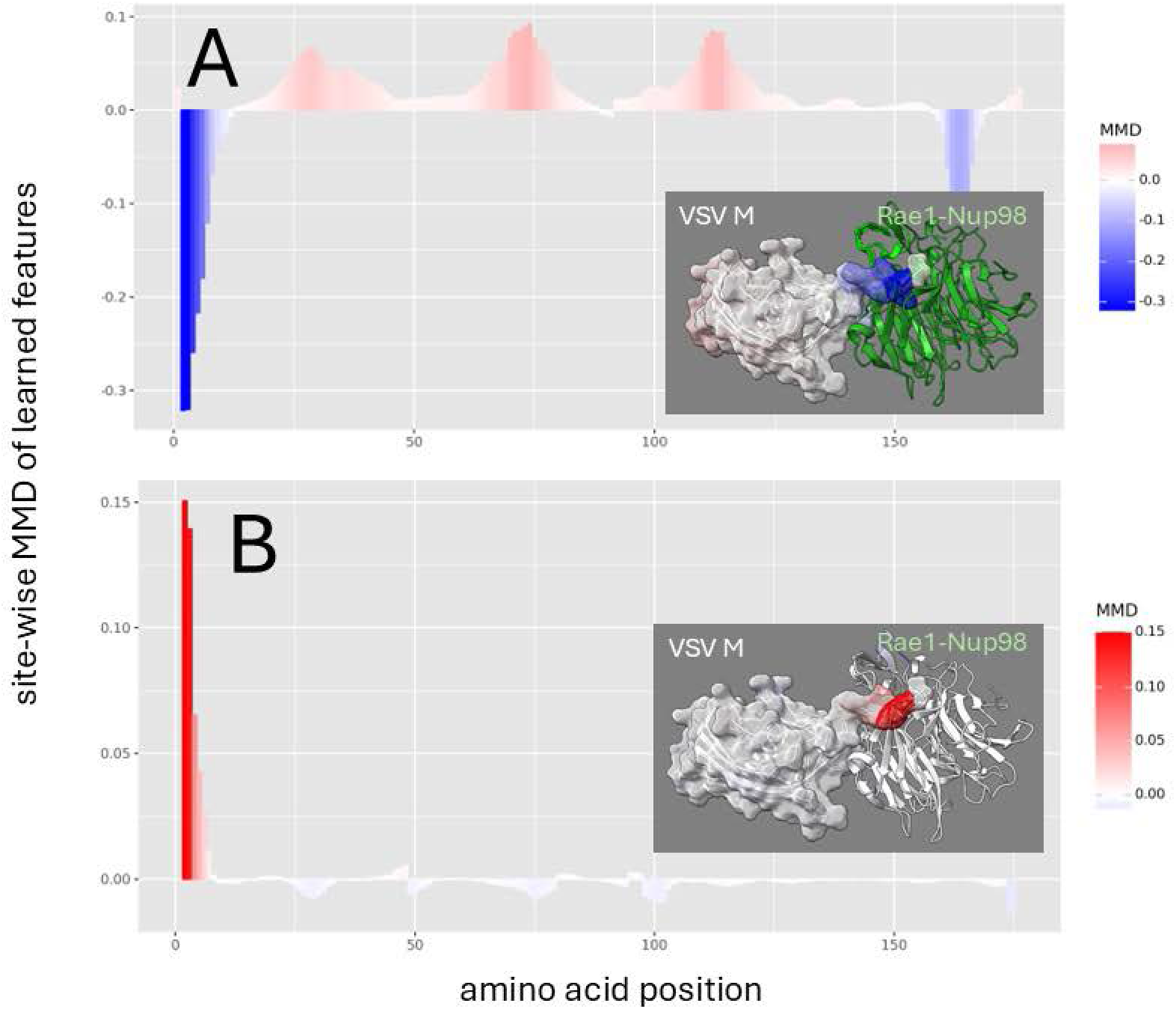
Site-wise denoised molecular dynamics comparisons of (A) Rae1-Nup98 bound vs. unbound VSV M protein and (B) wild-type vs. M51R mutant Rae1-Nup98 bound VSV M protein. The location of this site is a amino acid position 3 on our plot. The method of denoising utilizes the maximum mean discrepancy (MMD) in learned features of local atom fluctuations that best distinguish the two dynamic states being compared (Babbit et al. 2024). The machine learning method employed is a Gaussian process kernel employing a simple radial basis function. Both comparisons highlight the singular importance of the non-random changes in dynamics created by (A) the presence of the functional methionine in the N-terminal region of VSV M protein during binding and (B) its replacement by arginine. As in Figure 2 the MMD is signed negative (blue) to indicate dampened motion and positive (red) to indicate amplified motion.

In KSHV ORF 10, we observe a simple role of the critical methionine in binding that is similar to its role in VSV M (Figure 4A) but a more complex indication of its role in mutational destabilization, where the much longer polypeptide extension used by KSHV ORF 10 still shows some sites still involved in Rae1-Nup98 interaction despite the mutation (Figure 4B). Unlike VSV M where the critical methionine in at the N terminus of the M protein, this dampening effect localized to the critical methionine is positioned near the C terminus of the ORF 10 protein. While introducing the M413R mutation in ORF10 results in a substantial shift in dynamic interactions (Figure 2). In contrast to the total reversal observed in VSV, where the entire finger-like region experienced amplified motion, KSHV exhibits a more nuanced response. While most of the finger-like region undergoes increased mobility, the furthest terminal segment remains relatively dampened near the original site of methionine, indicating that other sites may be involved in its interference with host mRNA transport.

**Figure 4.**
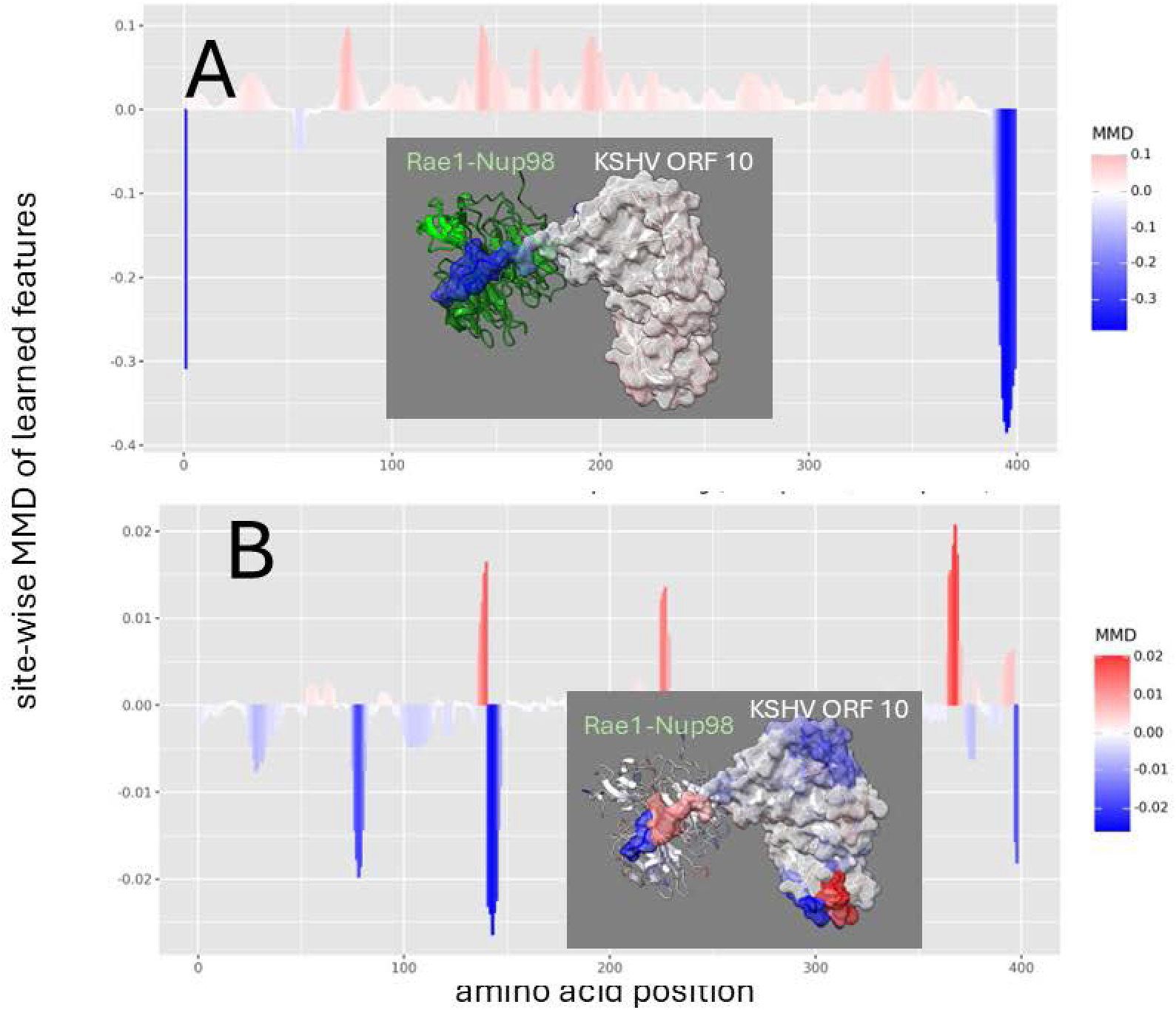
Site-wise denoised molecular dynamics comparisons of (A) Rae1-Nup98 bound vs. unbound KSHV ORF 10 protein and (B) wild-type vs. M413R mutant Rae1-Nup98 bound KSHV ORF 10 protein. The location of this mutation is shown at position 396 on our plots. The method of denoising utilizes the maximum mean discrepancy (MMD) in learned features of local atom fluctuations that best distinguish the two dynamic states being compared (Babbit et al. 2024). The machine learning method employed is a Gaussian process kernel employing a simple radial basis function. Both comparisons highlight the singular importance of the non-random changes in dynamics created by (A) the presence of the functional methionine in the C-terminal region of KSHV ORF 10 protein during binding and (B) its replacement by arginine. As in Figure 2 the MMD is signed negative (blue) to indicate dampened motion and positive (red) to indicate amplified motion.

The denoised dynamics comparative analysis of the SARS-CoV-1 ORF6-Rae1-Nup98 complex dynamics is depicted in Figure 5. Given that SARS-CoV-1 ORF6 is a particularly short sequence, interpreting its dynamic changes poses challenges, as observed fluctuations may stem from its fragmentary nature rather than binding or mutation effects. While the overall thermodynamic effect is clear (Figure 2), the sites identified by the MMD calculations show non-random dampening is observed primarily at the terminal ends of ORF6 rather than at the critical methionine site. This pattern likely reflects the untethered nature of the terminal regions rather than a direct binding-induced stabilization. A comparison of SARS-CoV-1 and SARS-Cov-2 ORF6 (Supplemental Figures 4-5) show that there may be subtle differences in how ORF6 mutations drive changes in functional dynamics of the host target complex. In SARS-CoV-1, a functional loop in Nup98 appears to be more stabilized while interacting with a less stabilized ORF 6. However, in SARS-CoV-2, Nup 98 appears destabilized while the dynamics of ORF 6 are unchanged. It is likely that these differences may simply reflect different temporal snapshots of a more complex process common to both.

**Figure 5.**
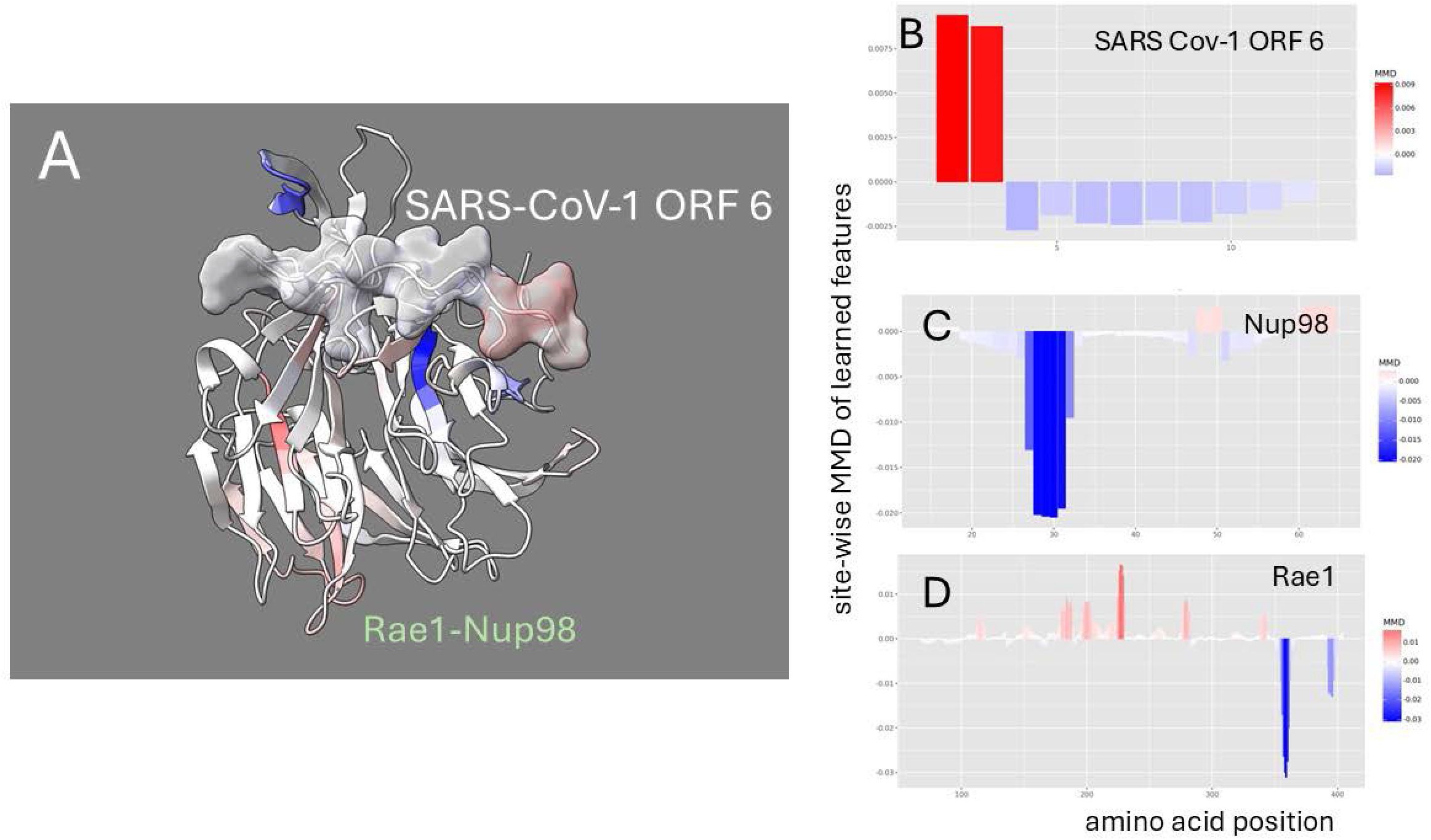
Site-wise denoised molecular dynamics comparisons of (A) Rae1-Nup98 bound vs. unbound SARS-CoV-1 ORF 6 protein and (B) wild-type vs. M to R mutant Rae1-Nup98 bound SARS-CoV-1 ORF 6 protein. The method of denoising utilizes the maximum mean discrepancy (MMD) in learned features of local atom fluctuations that best distinguish the two dynamic states being compared (Babbit et al. 2024). The machine learning method employed is a Gaussian process kernel employing a simple radial basis function. Both comparisons highlight the singular importance of the non-random changes in dynamics created by (A) the presence of the functional methionine in the center region of SARS-CoV-1 ORF 6 protein during binding and (B) its replacement by arginine. As in Figure 2 the MMD is signed negative (blue) to indicate dampened motion and positive (red) to indicate amplified motion.

While it is generally presented in the literature that these critical methionine residues are functionally conserved across viral taxa, we show that the alignment shows that acidic flanking regions are more variable (Supplemental Figure 6). Given the large evolutionary distance, the major structural differences between VSV M and KSHV ORF10 (Figure 1), and the rapid protein-level evolution that typically occur in viruses, we suggest that a history of convergent evolution could be equally supported by the sequence alignment. Further work with a phylogenetically broader scope along with ancestral sequence reconstruction could further illuminate the history of these common functional methionine containing extension regions.

## Discussion

In this work, by comparing the molecular dynamics of nucleoporin bound vs. unbound viral proteins, we have documented a common functional mechanism for the inhibition of mRNA export among three structurally and phylogenetically diverse types of virus. All of these involve a methionine strategically placed within acidic or hydrophilic terminal regions of non-homologous viral proteins, strongly suggesting a process of convergent viral evolution among potentially many different pathogens in how they combat host defenses during infection.

Our method leveraged explicit solvent GPU accelerated molecular dynamics simulations that provide single-atom resolution machine learning feature data towards the calculation of comparative discrepancies between learned features (i.e. maximum mean discrepancy derived from a Gaussian Process learner) that were directly related to the binding of viral proteins to the Rae1-Nup98 complexes in the host nucleopore. This method cheaply and efficiently provides information about molecular functions at a single site resolution that is often beyond what can be done easily in a wet lab setting. It also allows for analyses with broad taxonomic scope, which is also difficult where most labs are set up to do very specialized work within singe groups of viruses. While our computational evidence supports the role of methionine in viral-host interactions, limitations exist. The accuracy of our models depends on the resolution of available structural data as well as common assumptions that come with the underlying protein force fields applied to the system, which may not fully account for external factors like post-translational modifications, environmental conditions, or alternative binding partners. And, as mentioned above, comparative molecular dynamics insights for SARS-CoV were limited due to the small size of ORF6.

Disrupting viral mRNA nuclear export is an emerging antiviral strategy aimed at interrupting a critical step in the viral infectious cycle. Preventing this process can suppress viral replication and restore host antiviral gene expression. One approach involves inhibition of the host export receptor CRM1 (also known as XPO1 or Exportin 1), which is exploited by viruses such as HIV-1, influenza A, and herpes simplex virus type 1 to export unspliced or incompletely processed viral RNAs (26, 27). Leptomycin B (LMB), a potent CRM1 inhibitor, prevents nuclear export by covalently modifying a critical cysteine residue in CRM1 (28, 29). Selinexor, a selective CRM1 inhibitor is approved for treating multiple myeloma (30) and relapsed or refractory diffuse large B-cell lymphoma (DLBCL) in humans (31) and has shown antiviral activity against SARS-Cov-2 (32); suggesting potential for repurposing. Two recent studies have shown that 4-Octyl itaconate (4-OI), a cell-permeable derivative of the endogenous metabolite itaconate, suppresses replication of multiple influenza A virus strains by targeting CRM1, thereby blocking the nuclear export of viral ribonucleoproteins (33, 34).

A complementary strategy involves targeting viral proteins that hijack the mRNA export machinery. For example, therapeutics that disrupt the ability of the VSV M, SARS-CoV ORF6, or KSHV ORF10 protein’s ability to interact with the Rae1–Nup98 complex could selectively impair viral mRNA export and gene expression while sparing host functions. Blocking viral mRNA export can also restore host antiviral responses, including the production of interferon-stimulated genes (ISGs). Viral interference with Rae1–Nup98 can suppress ISG mRNA export and protein production (2, 3, 14, 15, 17, 35). Thus, targeting this axis not only inhibits viral gene expression but also amplifies innate immune signaling.

Finally, mRNA-based therapies offer an indirect but promising modality. These therapies can be engineered to deliver antiviral effector proteins, immune-stimulatory cytokines, or dominant-negative inhibitors of viral proteins, thereby enhancing immune recognition and suppressing replication (36, 37). Together, these strategies demonstrate that targeting viral mRNA nuclear export—either by inhibiting host export factors, disrupting viral-host interactions, or enhancing immune responses—represents a viable avenue for antiviral intervention, particularly against viruses that manipulate the nuclear pore complex for replication.

In summary, the application of comparative molecular dynamics simulations with **ATOMDANCE** software provides critical insight into dynamic molecular fluctuations that are often inaccessible by experimental methods alone. By integrating high-resolution structural modeling with future experimental validation, this approach may offer a powerful framework for dissecting viral–host interactions and pave the way for deeper mechanistic understanding of viral pathogenesis, while informing the rational design of targeted antiviral strategies.

Future studies could leverage ATOMDANCE to simulate interactions between the complex and known selectively expressed ssRNA transcripts to test our proposed mechanism. As a possible case of convergent evolution, we are currently investigating these shared patterns of host mRNA export inhibition, as this may represent a key mechanism that additional viruses exploit to suppress host antiviral responses. Despite the insights gained, further research is needed to fully characterize the role of Rae1-Nup98 in viral mRNA export inhibition. The interaction remains incompletely understood, particularly regarding KSHV’s selective inhibition mechanism.

## Acknowledgements

We thank Dr. Douglas Lyles for his support and insightful comments on the manuscript. We thank the College of Science at the Rochester Institute of Technology (RIT) for helping to support this work. We also thank the College of Science Summer Undergraduate Research Program and the RIT Honors Program for supporting one of the undergraduates who contributed to this work. This research was supported by NIH grant 1R15CA246419 (to M. C. F.). The funders had no role in study design, data collection and interpretation, or the decision to submit the work for publication.

## Figure Legends

**Supplemental Figure 1.**
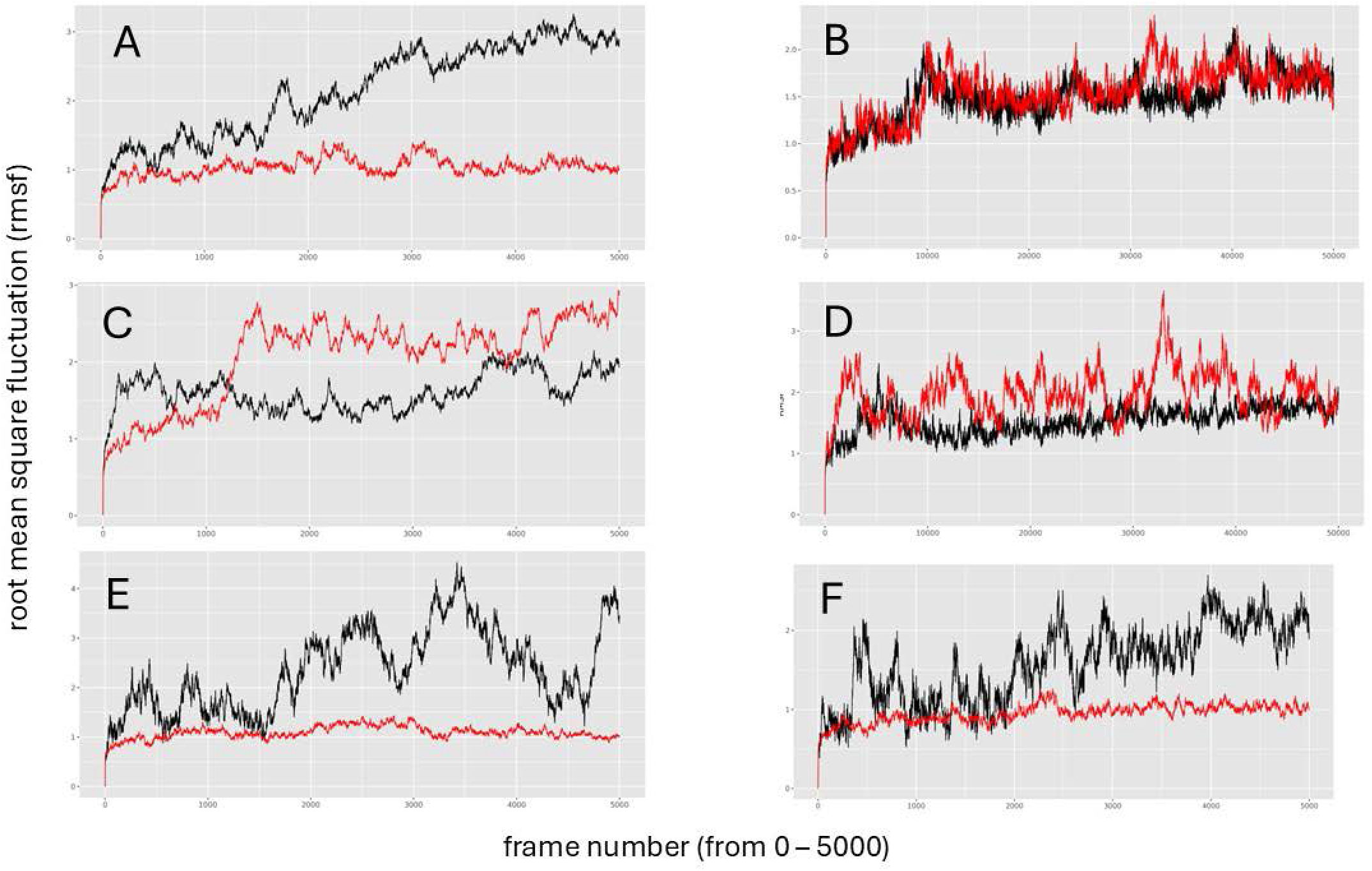
Root mean square fluctuation plots for the computational binding. experiments on (A-B) VSV M, (C-D) KSHV ORF 10, (E) SARS-CoV-1 ORF 6 and (F) SARS-Cov-2 ORF 6. The bound vs unbound rmsf values are shown resp as red and black and appear stable at both 1ns (A,C,E and F) and 10ns (B, D) time intervals.

**Supplemental Figure 2.**
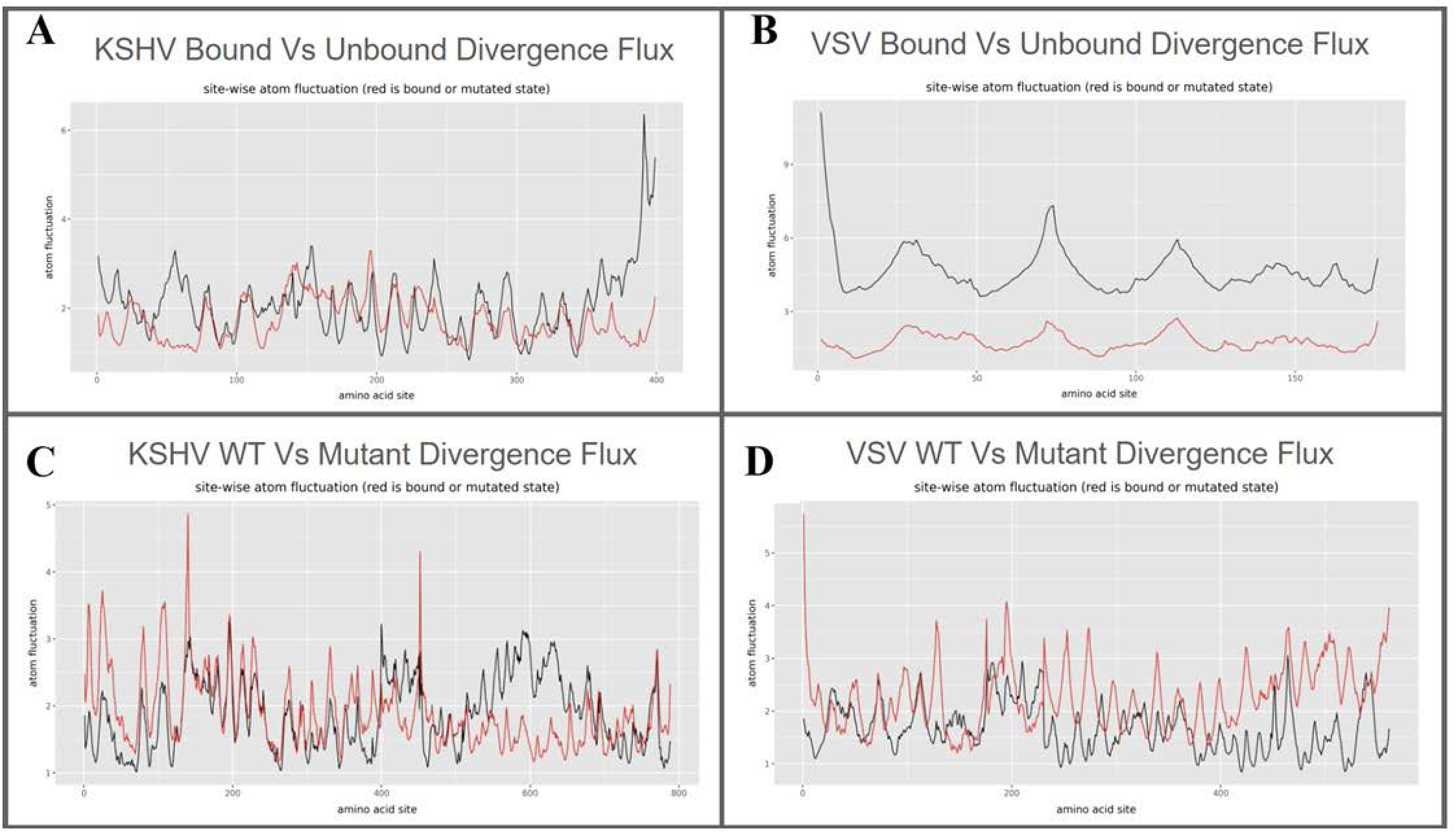
Site-wise average root mean square fluctuation plots comparing (A-B) bound vs unbound KSHV ORF 10 and VSV M structures and (C-D)

**Supplemental Figure 3.**
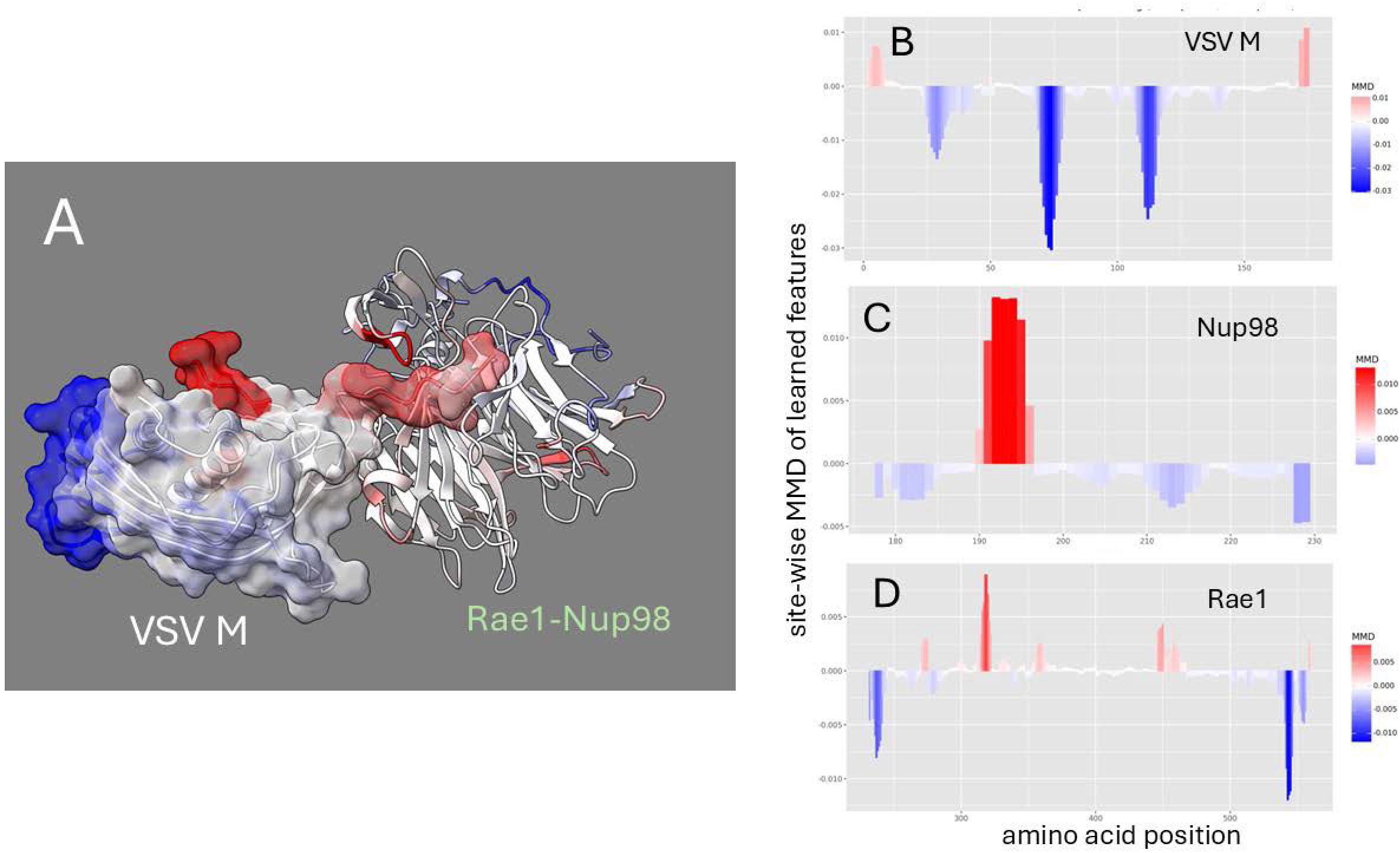
Site-wise denoised molecular dynamics comparisons. (A-B) wild-type vs. D52G mutant Rae1-Nup98 bound VSV M protein. Here is shown the additional amplification in functional sites of the host Rae1-Nup98 (C-D) as it tries to hold onto the mutant viral ORF 6 that is lacking a key acidic flanking residue in the region required for binding. The method of denoising utilizes the maximum mean discrepancy (MMD) in learned features of local atom fluctuations that best distinguish the two dynamic states being compared (Babbit et al. 2024). As in Figure 2 the MMD is signed negative (blue) to indicate dampened motion and positive (red) to indicate amplified motion.

**Supplemental Figure 4.**
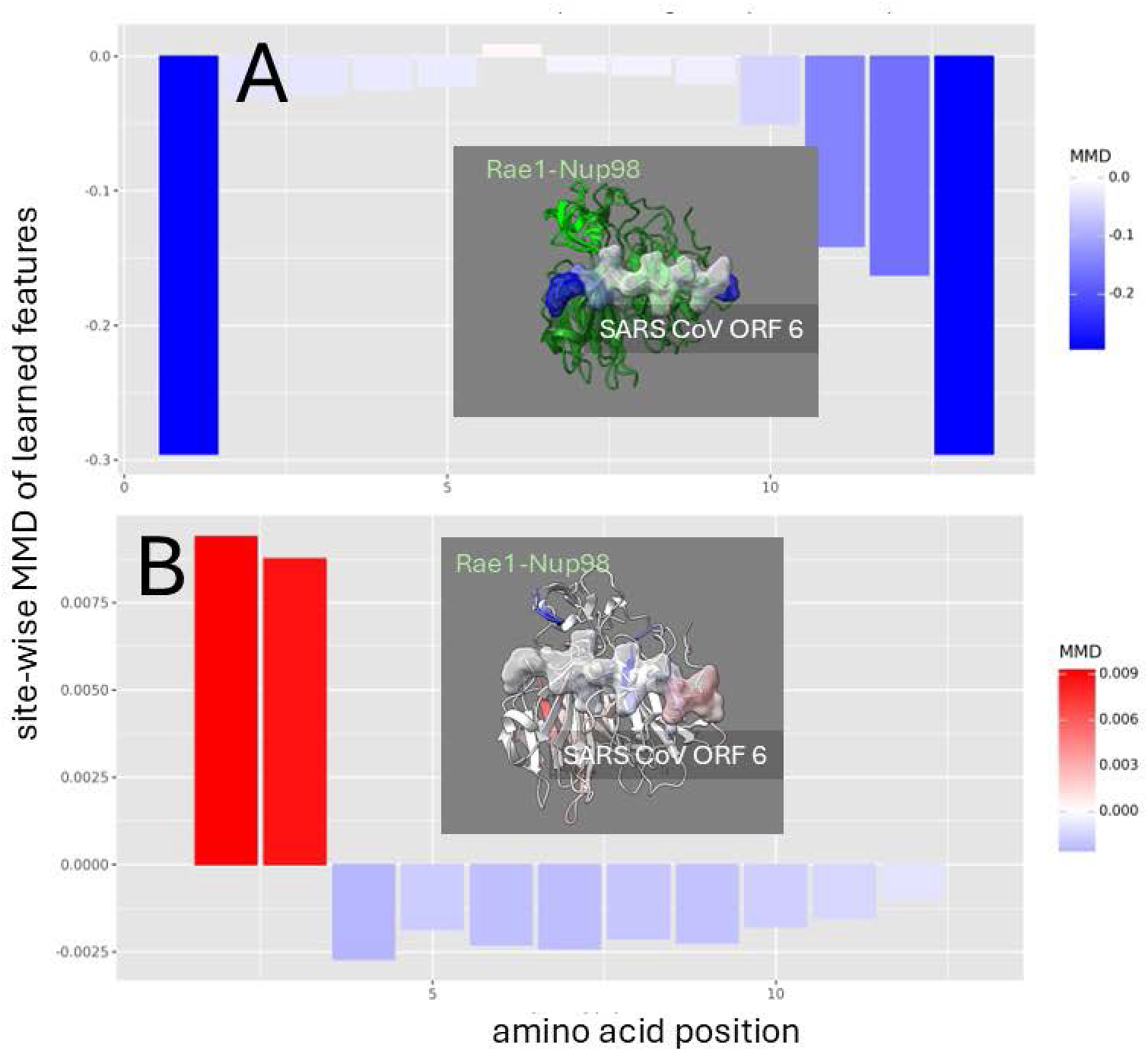
Site-wise denoised molecular dynamics comparisons. (A-B) wild-type vs. M to R mutant Rae1-Nup98 bound SARS-CoV-1 ORF 6 protein. Here is shown the additional dampening in functional sites of the host Rae1-Nup98 (C-D) as it tries to hold onto the mutant viral ORF 6 that is lacking the key methionine region required for binding. The method of denoising utilizes the maximum mean discrepancy (MMD) in learned features of local atom fluctuations that best distinguish the two dynamic states being compared (Babbit et al. 2024). As in Figure 2 the MMD is signed negative (blue) to indicate dampened motion and positive (red) to indicate amplified motion.

**Supplemental Figure 5.**
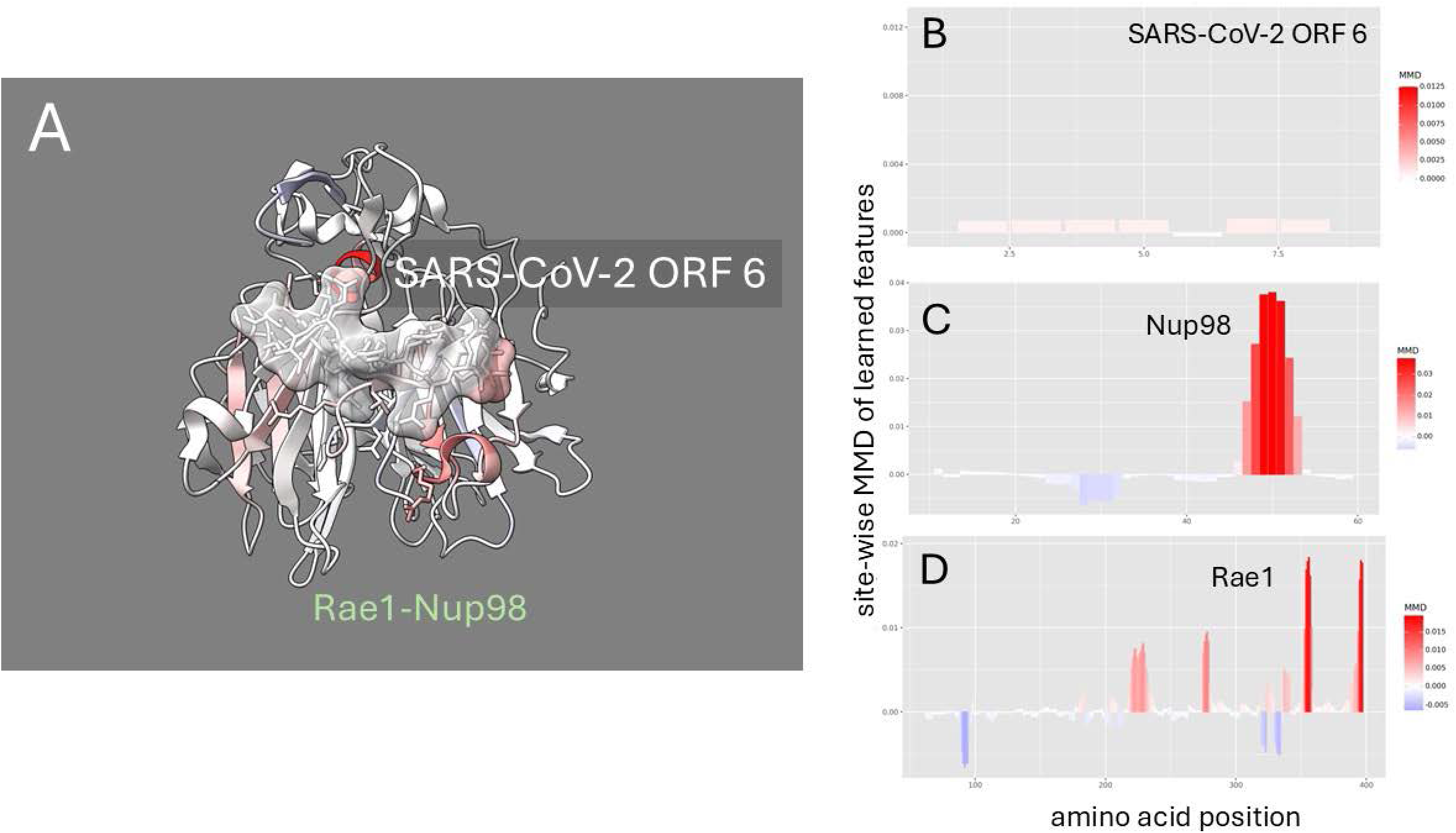
Site-wise denoised molecular dynamics comparisons. (A-B) wild-type vs. M to R mutant Rae1-Nup98 bound SARS-CoV-2 ORF 6 protein. Here is shown the additional amplification in functional sites of the host Rae1-Nup98 (C-D) as it tries to hold onto the mutant viral ORF 6 that is lacking the key methionine region required for binding. The method of denoising utilizes the maximum mean discrepancy (MMD) in learned features of local atom fluctuations that best distinguish the two dynamic states being compared (Babbit et al. 2024). As in Figure 2 the MMD is signed negative (blue) to indicate dampened motion and positive (red) to indicate amplified motion.

**Supplemental Figure 6.**
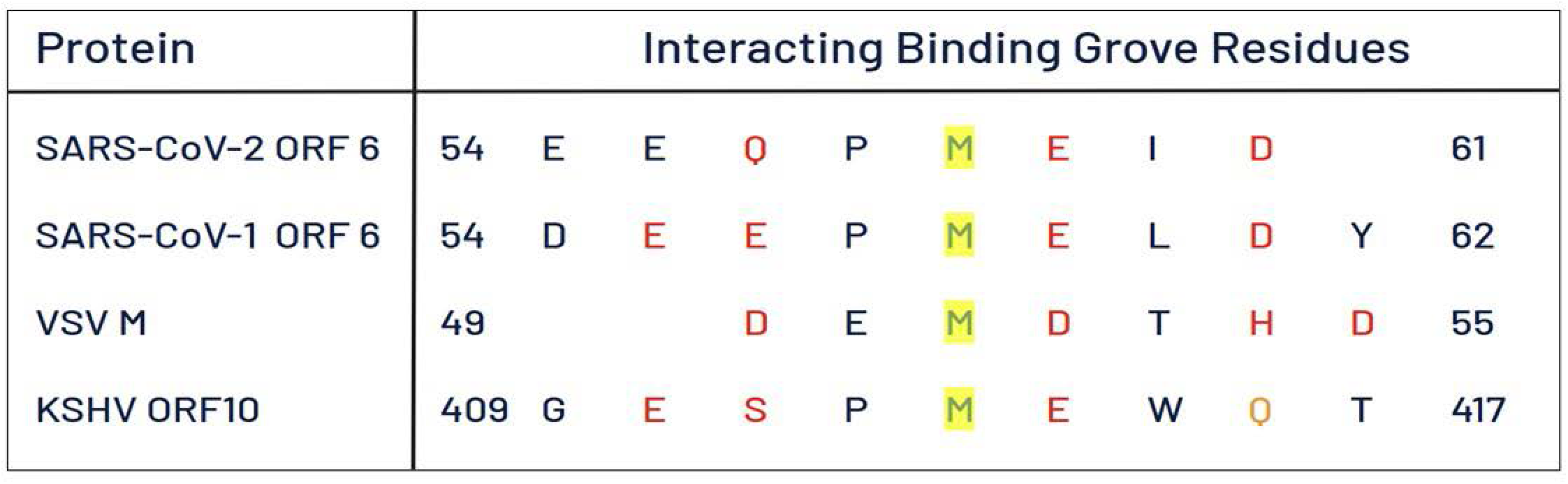
Comparative sequence alignment of the acidic flanked functional methionine (yellow). Acidic flanking residues are show in red. Although previous literature calls this methionine to be evolutionarily conserved, the high variability of the residues in the sites flanking the methionine is also suggestive that the functional methionine may be the result of convergent evolution of a similar mechanism among these highly divergent and rapidly evolving viral proteins.

## Notes

### Competing Interest Statement

The authors have declared no competing interest.

### Summary of Updates

No major cahnges have been made

